# The sperm protein Spaca6 is essential for fertilization in zebrafish

**DOI:** 10.1101/2021.11.19.469324

**Authors:** Mirjam I. Binner, Anna Kogan, Karin Panser, Alexander Schleiffer, Victoria E. Deneke, Andrea Pauli

**Affiliations:** Research Institute of Molecular Pathology (IMP), Vienna BioCenter (VBC), Campus-Vienna-Biocenter 1, 1030 Vienna, Austria

**Keywords:** fertilization, zebrafish, sperm-egg interaction, gamete, sperm, reproduction

## Abstract

Fertilization is a key process in all sexually reproducing species, yet the molecular mechanisms that underlie this event remain unclear. To date, only a few proteins have been shown to be essential for sperm-egg binding and fusion in mice, and only some are conserved across vertebrates. One of these conserved, testis-expressed factors is SPACA6, yet its function has not been investigated outside of mammals. Here we show that zebrafish *spaca6* encodes for a sperm membrane protein which is essential for fertilization. Zebrafish *spaca6* knockout males are sterile. Furthermore, Spaca6-deficient sperm have normal morphology, are motile, and can approach the egg, but fail to bind to the egg and therefore cannot complete fertilization. Interestingly, sperm lacking Spaca6 have decreased levels of another essential and conserved sperm fertility factor, Dcst2, revealing a previously unknown dependence of Dcst2 expression on Spaca6. Together, our results show that zebrafish Spaca6 regulates Dcst2 levels and is required for binding between the sperm membrane and the oolemma. This is in contrast to murine SPACA6, which was reported not to be required for sperm-egg membrane binding but necessary for fusion. These findings demonstrate that Spaca6 is essential for zebrafish fertilization and is a conserved sperm factor in vertebrate reproduction.

## Introduction

Fertilization is an essential process which ensures reproductive success and species survival. Multiple events, such as sperm migration through the female reproductive tract, sperm activation, gamete fusion and egg activation are necessary for fertility and contribute to the formation of a new organism (Bianchi and Wright, 2016; Stein *et al.*, 2020; Deneke and Pauli, 2021). Despite its critical role, little is known about the molecular mechanisms mediating fertilization, particularly sperm-egg interaction and fusion.

In recent years, several proteins have been shown to be essential for sperm-egg interaction in mammals. Most notably, mammalian testis-expressed IZUMO1 and its receptor JUNO on the egg membrane (oolemma) form a complex which is needed for binding of the two gametes prior to fusion (Inoue *et al.*, 2013; Bianchi and Wright, 2016). *Izumo1* knockout (KO) sperm can penetrate the zona pellucida (ZP), a glycoprotein coat surrounding the mammalian oocyte, but fail to fuse with the oolemma (Inoue *et al.*, 2005). CD9, an integral membrane protein expressed on the oolemma, has also been shown to be required for gamete fusion (Kaji *et al.*, 2000; le Naour *et al.*, 2000; Miyado *et al.*, 2000). Furthermore, other sperm factors – fertilization influencing membrane protein (FIMP), sperm-oocyte fusion required 1 (SOF1), transmembrane protein 95 (TMEM95), sperm acrosome associated 6 (SPACA6) and the DC-STAMP-like domain-containing proteins DCST1 and DCST2 – have been shown to be required for sperm-egg fusion in mice (Barbaux et al., 2020; Noda et al., 2020, 2021; Inoue et al., 2021). Despite the many factors shown to be necessary for sperm-egg fusion, the combination of these factors appears not to be sufficient to drive fusion in a heterologous system. Noda and colleagues expressed the essential sperm factors involved in fusion, including IZUMO1, in HEK293T cells and observed that HEK293T cells were able to bind but were unable to fuse to ZP-free eggs, indicating the possible need for additional molecules that are involved in this process (Noda *et al.*, 2020). Despite these recent advances, the molecular mechanisms of gamete fusion and the interplay between the various sperm and egg factors remain unclear.

Zebrafish has recently emerged as a model organism to study vertebrate fertilization due to its genetic tractability, access to a large number of gametes and external fertilization. The first factor that was discovered to be essential in zebrafish fertilization was Bouncer - a short three-finger-type protein anchored to the egg membrane (Herberg *et al.*, 2018). *Bouncer^−/−^* females are sterile and sperm is unable to bind to Bouncer-deficient eggs, suggesting that Bouncer is involved in sperm-egg membrane binding (Herberg *et al.*, 2018). Interestingly, the mammalian homolog of Bouncer is SPACA4, which is present in sperm and is involved in penetrating the zona pellucida (Fujihara *et al.*, 2021). Moreover, we have shown that DCST1 and DCST2 are also necessary for zebrafish fertilization (Noda *et al.*, 2021). However, while in mice *Dcst1* and *Dcst2* KO sperm can bind to, but rarely fuse with oocytes (Inoue et al., 2021; Noda et al., 2021), zebrafish *dcst1/2* KO sperm already show defects in sperm-egg binding (Noda *et al.*, 2021). Therefore, zebrafish represents an interesting model system for the study of fertilization; despite the presence of mammalian fertilization factors in fish (Dcst1/2, Bouncer/SPACA4), the currently characterized factors show functional divergence in the fertilization process (Fujihara *et al.*, 2021; Noda *et al.*, 2021). Elucidating the role of conserved fertilization factors in both mammalian and non-mammalian vertebrate species will help to uncover common themes as well as distinct functionalities. Here, we investigate the role of Spaca6 in fertilization in zebrafish.

## Results

### Spaca6 is a conserved testis-expressed membrane protein

To study the role of Spaca6 in zebrafish fertilization, we first identified the full-length *spaca6* mRNA and protein sequence in zebrafish. *Spaca6* has two gene annotations in the most recent zebrafish genome release (GRCz11): a predicted protein-coding gene containing 7 exons (NCBI: XM_021466914.1) and a predicted non-protein coding gene containing 8 exons (ENSEMBL: ENSDART00000155083.2). In addition to the difference in exon numbers, we noticed that the predicted NCBI protein-coding annotation lacked the N-terminus including the signal peptide, which is present in mammalian SPACA6 homologs (Noda *et al.*, 2020). To identify the sequence of the full-length *spaca6* transcript that is expressed in testis, we isolated RNA from wild-type zebrafish testes and amplified a 954-nt region from cDNA that contained an extended Spaca6 N-terminus and a total of 9 exons. Indeed, coverage tracks derived from RNA sequencing (Herberg *et al.*, 2018; Noda *et al.*, 2021) show expression peaks that align to the mapped transcript as well as specific expression of Spaca6 in testis (**Fig. 1A, Fig. S1A**), as has been previously reported in mammals (Barbaux *et al.*, 2020; Noda *et al.*, 2020).

**Figure 1.**
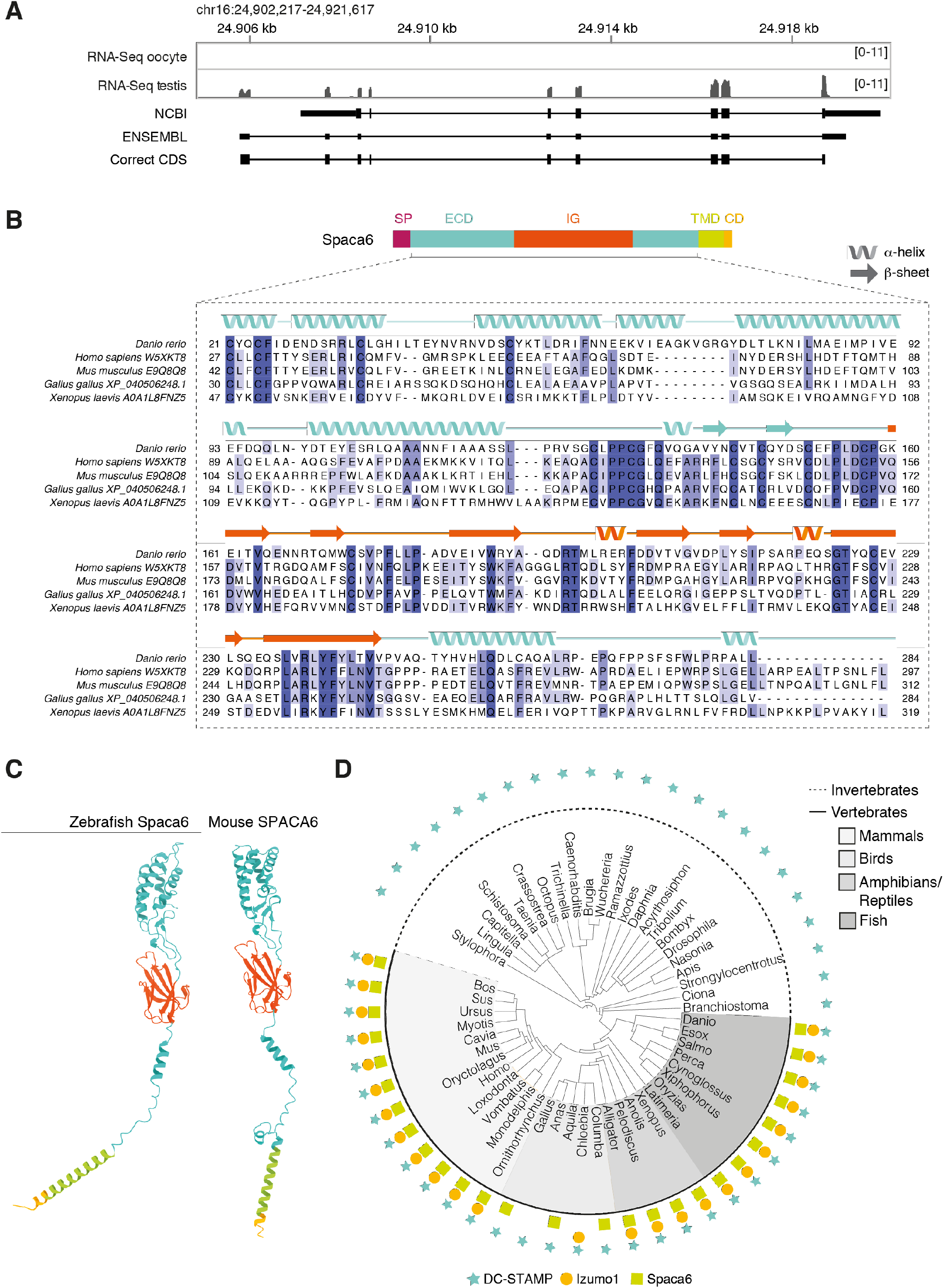
Identification of Spaca6 in zebrafish. **A.** Expression and genomic features of zebrafish *spaca6.* Coverage tracks for RNA sequencing data from oocyte and testis aligned with ENSEMBL (ENSDART00000155083.2) and NCBI (XM_021466914.1) annotations as well as the full-length, newly identified testis coding DNA sequence (correct CDS). Genomic coordinates are based on GRCz11. **B.** Protein domain structure of full-length Spaca6 and sequence alignment of the mature extracellular region of selected vertebrate Spaca6 proteins. Secondary structure prediction of the zebrafish Spaca6 protein sequence is depicted above the sequence alignment (coil, alpha-helix; arrow: beta-sheet). The extracellular domain (ECD) is shown in blue, while the Immunoglobulin-like domain (IG) is shown in orange. SP, signal peptide; TMD, Transmembrane domain; CD, Cytoplasmic domain. **C.** Tertiary structure predictions of mouse and zebrafish Spaca6. Protein folding predictions of mature Spaca6 (lacking the signal peptide sequence) were performed using AlphaFold2 (Jumper *et al.*, 2021). Extracellular domain, blue; Ig-like domain, orange; Transmembrane domain, green; Cytoplasmic domain, yellow. **D.** Taxonomic tree of DC-STAMP-like proteins, Izumo1 and Spaca6 across vertebrates and invertebrates. DC-STAMP-like proteins (blue star) are conserved both in vertebrates and invertebrates; Izumo1 (orange circle) and Spaca6 (green square) are conserved only in vertebrates.

The full-length zebrafish *spaca6* transcript encodes for a 318-amino acid single-pass membrane protein that resembles mammalian SPACA6 proteins in their sequence and tertiary structure (**Fig. 1B-C**). It contains an N-terminal signal peptide, followed by an extracellular region containing an alpha-helical domain (extracellular domain, ECD) and an immunoglobulin (Ig) fold domain (IG), a transmembrane domain and a short cytoplasmic domain (**Fig. 1B**). Alignment of Spaca6 protein sequences from different vertebrates shows that Spaca6 is conserved in its amino acid sequence and secondary structure elements (**Fig. 1B**).

Similar to mouse SPACA6, whose extracellular region has been shown to closely resemble that of IZUMO1 (Nishimura *et al.*, 2016), Alphafold2 (Jumper *et al.*, 2021) predicts the extracellular domain of zebrafish Spaca6 to fold into a four-helix bundle followed by an Ig-like domain comprised of beta sheets (**Fig. 1C**). In order to taxonomically map Spaca6 homologues, we performed iterative BLASTP searches. This revealed that - similar to Izumo1 - Spaca6 homologs are present across vertebrates, while proteins containing a DC-STAMP-like domain, such as Dcst1 and Dcst2, show a much broader distribution across both vertebrates and invertebrates (**Fig. 1D**). Together, the conservation of Spaca6 across vertebrates and predicted similarities in tertiary structure with the mammalian homologs motivated us to assess the functional role of Spaca6 in zebrafish.

### Spaca6 is essential for male fertility in zebrafish

Using CRISPR/Cas9-mediated mutagenesis, we generated zebrafish lacking Spaca6 protein. To this end, gRNAs were designed to target the 3^rd^ and 4^th^ exons of the full-length *spaca6* gene. A mutant harboring an 86-bp deletion was recovered, which resulted in the introduction of a premature stop codon **(Fig. 2A, Fig. S1B-C)**. To test whether Spaca6 is required for fertilization in zebrafish, we assessed the fertilization rates of wild-type, heterozygous and homozygous KO fish (**Fig. 2B-C**). While *spaca6^+/−^* males had fertilization rates comparable to wild type, *spaca6^−/−^* males were sterile (**Fig. 2C**). Consistent with the expression of Spaca6 exclusively in the male germline (**Fig. S1A**), *spaca6^−/−^* females were fertile (**Fig. 2C**), revealing that Spaca6 is only required for male fertility. To confirm that the observed fertilization defect of *spaca6^−/−^* males was caused by the absence of Spaca6 protein, we generated a rescue line that ubiquitously expresses Spaca6 in a *spaca6^−/−^* background *(spaca6^−^ tg[actb2::spaca6-T2A-GFP]),* hereafter referred to as *spaca6^−/−^; tg(spaca6).* Transgenic Spaca6 partially rescued the fertilization defect of *spaca6* KO males (**Fig. 2C**), suggesting that the fertility defect is indeed due to a lack of Spaca6.

**Figure 2.**
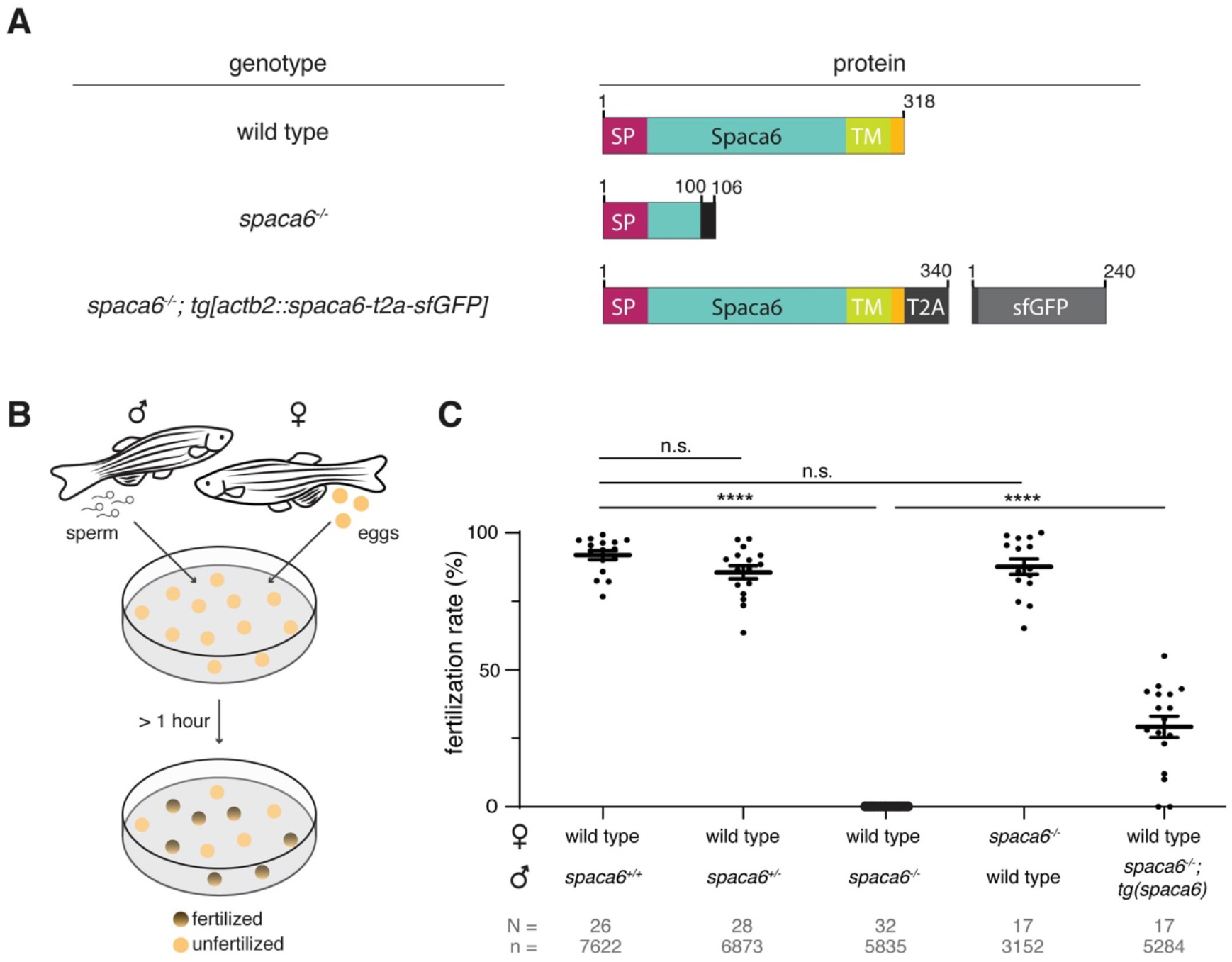
Zebrafish Spaca6 is essential for male fertility. **A.** Diagrams of Spaca6 proteins made in wild-type, *spaca6^−/−^* and rescue *(spaca6^−/−^; tg[actb2::spaca6-t2a-GFP])* zebrafish lines. An 86-nt deletion in the *spaca6^−/−^* line leads to a retained sequence from intron 2 and a premature stop-codon, resulting in a dysfunctional truncated protein (translated part of the intron is shown in black). In the rescue line, a SGGSG spacer and a part of the viral T2A protein result in a C-terminal 23-aa addition. The amino acid position is given above the scheme. Signal peptide (SP), red; Extracellular domain, blue; Transmembrane domain (TM), green; Cytoplasmic domain, yellow. **B.** Scheme of a mating assay performed to quantify fertilization rates. Fish and gametes are not drawn to scale. **C.** Quantification of fertilization rates. Fertilization rates were calculated by counting the number of embryos that developed beyond the one-cell stage. ****, p < 0.0001 (Mann-Whitney test); n.s., not significant; error bars, standard deviation (SD); N = number of crosses; n = number of embryos.

### Spaca6 KO sperm is motile but fails to bind to the egg membrane

To functionally dissect the fertilization defect in *spaca6^−/−^* males, we first assessed whether mutant sperm were morphologically normal. Differential interference contrast (DIC) images did not show any gross morphological differences between *spaca6^−/−^* and wild-type sperm **(Fig. 3A)**. Furthermore, *spaca6^/^* and wild-type sperm had a similar sperm head size and tail length (**Fig. 3B**). In addition, we reasoned that another parameter that may affect sperm fertilizing ability is sperm motility. However, both wild-type and mutant sperm showed comparable motility and directed displacement (**Fig. 3C-D**). Therefore, the fertilization defect of *spaca6* KO sperm is neither due to morphological defects nor due to an inability of sperm to undergo activation and directed movement.

**Figure 3.**
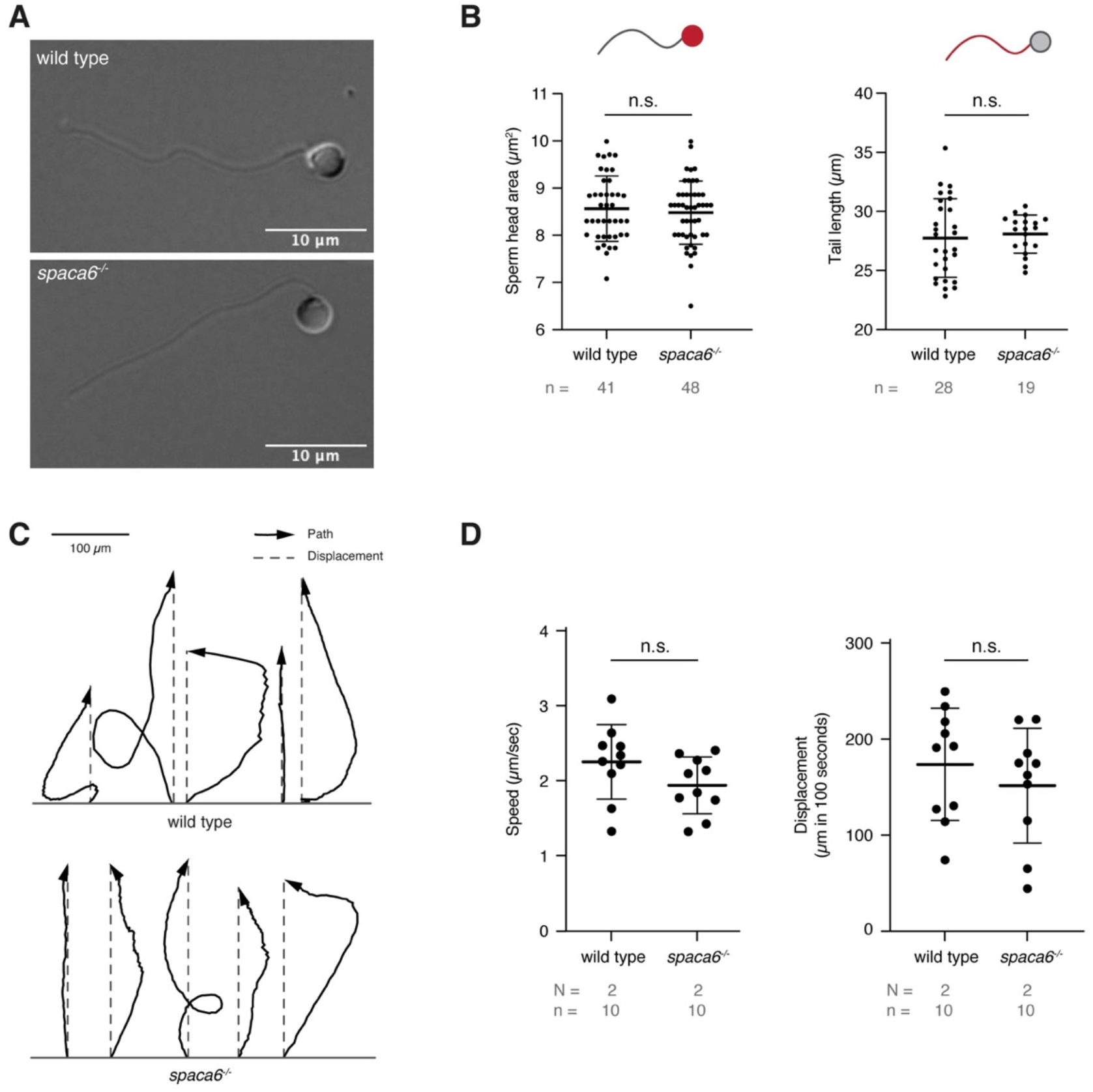
*Spaca6* KO sperm is morphologically normal and motile. **A.** Representative differential interference contrast (DIC) images of wild-type and *spaca6^−/−^* sperm. Scale bar = 10 μm. **B.** Measurements of the head area (left) and tail length (right) of wild-type and *spaca6^−/−^* sperm. n.s., not significant; error bars, SD; n = number of sperm. **C.** Representative tracks of wild-type and *spaca6^−/−^* sperm. Tracks (black arrows) were rotated and aligned at their origin. The relative displacement (grey dashed line) is overlaid. **D.** Sperm speed and displacement (per 100 sec). n.s., not significant; error bars, SD; N = number of independent experiments; n = number of sperm.

We next tested whether *spaca6* KO sperm is able to approach the micropyle, a funnel-shaped structure within the egg coat that serves as the single sperm entry point in fish. Wild-type and *spaca6^−/−^* sperm were added to wild-type eggs and a time-lapse of sperm approaching the micropyle region was recorded. Both wild-type and mutant sperm were able to reach the micropyle within the same time-frame (**Fig. 4A, Movie 1**). However, while wild-type sperm remained in the micropyle, *spaca6^−/−^* sperm drifted away after a few minutes (**Fig. 4A, Movie 1**). This provided a first indication that wild-type sperm is able to stably adhere to the egg plasma membrane, while *spaca6^−/−^* sperm may fail to stably bind the egg surface.

**Figure 4.**
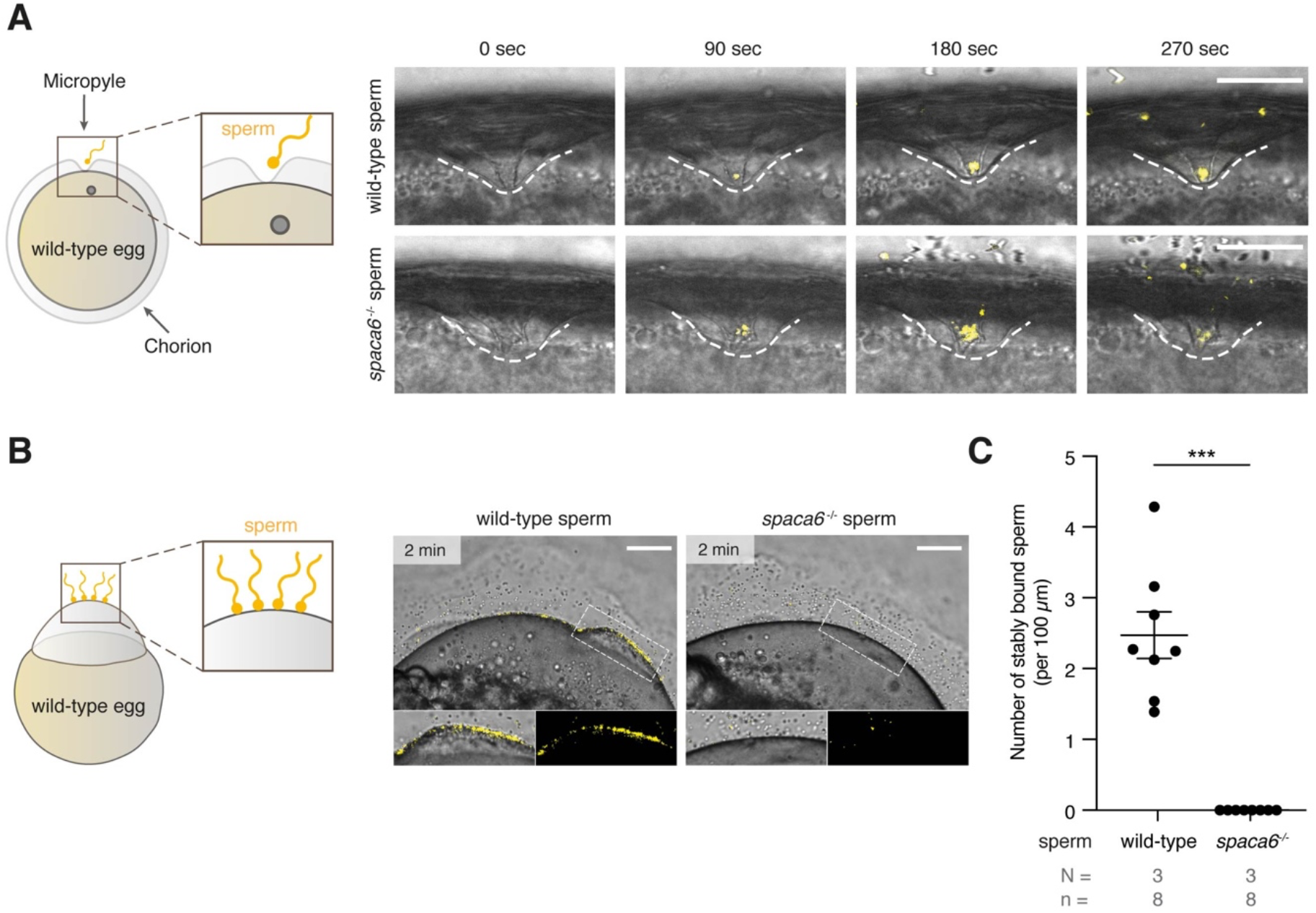
Zebrafish Spaca6 is essential for sperm-egg binding. **A.** Time-lapse images of sperm approaching the micropyle of wild-type eggs. Left: Experimental setup. Right: Representative time series of MitoTracker-labeled wild-type (top) and *spaca6^−/−^* (bottom) sperm (yellow) approaching the micropyle (white dashed lines). Scale bar = 50 μm. **B.** Sperm binding assay. Left: Experimental setup. Wild-type and *spaca6^−/−^* sperm were labeled with MitoTracker (yellow) and subsequently incubated with activated and dechorionated wild-type eggs. Right: Representative images of wildtype (top) and *spaca6^−/−^* (bottom) sperm binding to the surface of the egg 2 minutes after sperm addition. The boxed region is shown at higher magnification below. Scale bar = 100 μm. **C.** Quantification of sperm binding. Binding of sperm to the egg was quantified by assessing the number of stably bound sperm per 100 μm over a period of 2 minutes. Wild-type sperm were frequently found bound to the oolemma. ***, p < 0.001 (Mann-Whitney test); error bars, SD; N = number of independent experiments; n = number of sperm.

To directly test whether *spaca6^−/−^* sperm is defective in egg binding, we performed a sperm binding assay that we had previously established (Herberg *et al.*, 2018; Noda *et al.*, 2021). In short, wild-type eggs were activated with water and manually dechorionated to expose the entire egg surface. Sperm was then added and sperm binding rates were recorded via timelapse imaging. As observed in the sperm approach assay, *spaca6^−/−^* sperm was able to reach the egg, but failed to stably bind to the egg surface (**Fig. 4B, Movie 2**). This was in contrast to wild-type sperm, which was able to stably bind to the egg surface for at least one minute (**Fig. 4C**, **Movie 2**). Therefore, we conclude that Spaca6 is required for sperm to stably adhere to the egg plasma membrane in zebrafish.

### Dcst2 levels are reduced in spaca6 KO sperm

Recent work in mice has shown that SPACA6 levels are decreased in sperm lacking IZUMO1, DCST1 and/or DCST2 (Inoue et al., 2021), suggesting that these essential fertility factors may co-regulate each other. To investigate whether this regulation holds true reciprocally, we measured Dcst2 protein levels and localization in *spaca6^−/−^* sperm, using an antibody recognizing zebrafish Dcst2 (Noda *et al.*, 2021). Dcst2 levels were significantly decreased in *spaca6^−/−^* sperm, as revealed by immunoblotting (**Fig. 5A-B, Fig. S2**) and immunofluorescence imaging, which showed a reduction of Dcst2-positive foci around the sperm plasma membrane (**Fig. 5C**). The decrease in Dcst2 levels was partially restored in sperm of the *spaca6^−/−^; tg(spaca6)* rescue line, which is consistent with the partial rescue of fertility (**Fig. 2B**). Our data therefore suggests that Spaca6 regulates Dcst2 protein levels.

**Figure 5.**
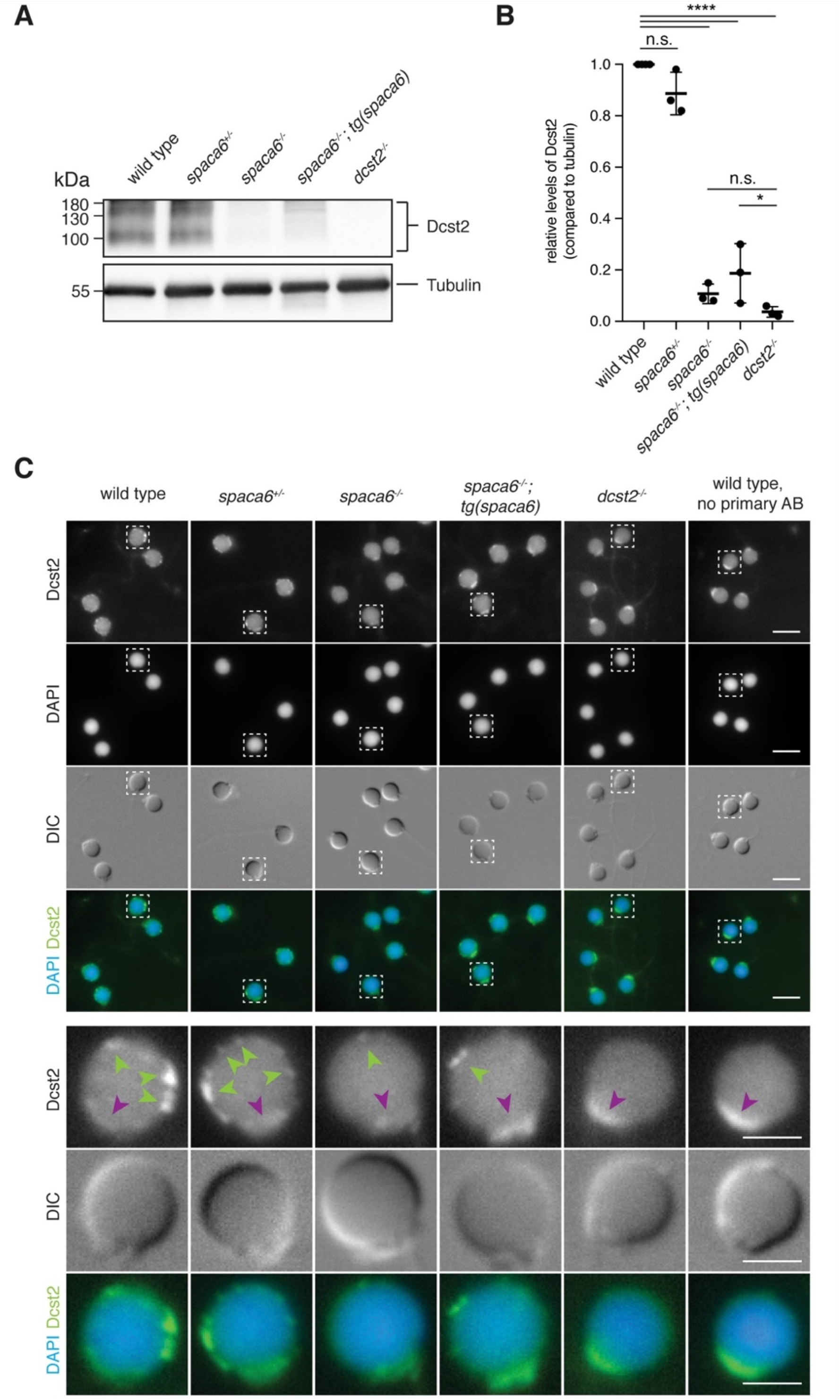
Dcst2 levels are reduced in zebrafish *spaca6* KO sperm. **A.** Immunoblot of sperm samples of different genotypes probed with antibodies against zebrafish Dcst2. Zebrafish Dcst2 is predicted to be a glycoprotein, resulting in a wider area of detected signal. Tubulin protein levels of the same blot are shown as loading control. The uncropped immunoblot is shown in Fig. S2. **B.** Quantification of sperm Dcst2 levels of different genotypes based on three independent immunoblots. Values were normalized to wild-type levels. Statistical significance was calculated using one-way ANOVA and multiple comparisons analysis: n.s., not significant (p > 0.05; ****, p < 0.0001; *, p = 0.045). **C.** Immunofluorescent images of Dcst2 protein in sperm heads of different genotypes. Upper panel: Overview images (scale bar = 5 μm). Lower panel: Higher magnification images of an individual exemplary sperm head (boxed in the upper panel) for each genotype (scale bar = 2 μm). Sperm were stained with DAPI (blue) to visualize nuclei and antibodies against zebrafish Dcst2 (green). Dcst2 localizes to distinct foci around the sperm head membrane (green arrowheads in the lower panel) in wild-type, *spaca6^+/−^* and *tg(spaca6)-rescued* sperm. Autofluorescence of the sperm midpiece appears as a uniform signal in the Dcst2 channel (magenta arrowhead).

## Discussion

Our results show that testis-expressed Spaca6 is essential for zebrafish fertilization. We make two major findings: (1) In zebrafish, the absence of Spaca6 leads to a disruption in the sperm’s binding ability to the egg surface and (2) zebrafish Spaca6 regulates the levels of another key fertilization factor, Dcst2.

Our study shows that Spaca6 has an essential role in fertilization outside of mammals. Even though Spaca6 is conserved among vertebrates (**Fig. 1B-D**), the step at which fertilization stalls in the absence of Spaca6 differs between zebrafish and mice. Mouse *Spaca6* KO sperm can penetrate the zona pellucida and bind to the oolemma, but fail to fuse with the egg (Barbaux *et al.*, 2020; Noda *et al.*, 2020). In contrast, our findings in zebrafish reveal that zebrafish Spaca6 is already required in a step before fusion can take place: sperm-egg membrane binding.

On a molecular level the observed difference could possibly be reconciled from mouse experiments involving IZUMO1. One possibility is that mouse *Spaca6* KO sperm is still able to bind to the oocyte due to the presence of IZUMO1, thereby providing a redundant role in sperm adhesion to the egg. In support of this notion, IZUMO1 protein levels as well as IZUMO1 relocation during the acrosome reaction were reported to be normal in *Spaca6* KO sperm (Barbaux et al., 2020; Inoue et al., 2021). Moreover, IZUMO1 was shown to be sufficient for cells to bind to the oocyte (Noda *et al.*, 2020), suggesting that one major role of IZUMO1 is to enable sperm to adhere to the egg surface. Although zebrafish have an Izumo1 ortholog (**Fig. 1C**), its mammalian binding partner on the egg surface, JUNO, appears to be absent in fish (Grayson, 2015). Izumo1’s presence may therefore not be sufficient to mediate binding between the two gametes in fish. Future experiments exploring the role of Izumo1 in zebrafish fertilization as well as potential co-regulation of Spaca6 and Izumo1 will help elucidate the molecular mechanisms in fish and mammalian systems alike.

Direct evidence that essential sperm factors co-regulate each other was recently reported in mice (Inoue et al., 2021). Inoue and colleagues showed that sperm lacking IZUMO1, DCST1 and/or DCST2 had undetectable levels of SPACA6 protein, which suggested that SPACA6 levels are dependent on the presence of each of these factors. Interestingly, SPACA6 protein levels were restored when IZUMO1 was transgenically expressed in *Izumo1* KO sperm, but not when DCST1/2 were expressed in *Dcst1/2* KO sperm, even though fertilization was rescued. One possibility is that SPACA6 levels were undetectable in the rescue but high enough for fertilization to be restored. Another possibility is that the role of SPACA6 can be by-passed by transgenic expression of DCST1/2, suggesting an indirect role of SPACA6 in fertilization. Our data shows that in zebrafish, transgenic expression of Spaca6 results in a partial rescue of fertilization (**Fig. 2C**), which is correlated with a partial restoration of Dcst2 levels (**Fig. 5**). However, it is currently unclear whether this partial rescue in fertilization is due to insufficient levels of Spaca6, Dcst1/2 or both. Together, this study and studies in mice point towards a coregulation of SPACA6 and DCST1/2. Inoue et al. demonstrated that DCST1/2 regulates the levels of SPACA6 (Inoue et al., 2021) and we show that zebrafish Spaca6 regulates Dcst2 levels (**Fig. 5**). Whether these factors act purely as stabilization factors or whether they also play a role in sperm-egg interaction remains to be tested. Further studies characterizing the molecular co-regulation of these factors will be necessary to understand their role in protein stability, binding and fusion.

In conclusion, Spaca6 is a conserved factor essential for fertilization in vertebrates, but its molecular function still remains unclear. Judging from the current data, there are several possible explanations to reconcile the role of Spaca6. Since Dcst2 levels are disrupted in *spaca6* KO sperm in zebrafish, Spaca6 may serve as a stabilization factor. While this idea has not yet been tested experimentally in mammals, loss of DCST1/2 protein levels in mammalian *Spaca6* KO sperm would provide direct evidence for a conserved role of Spaca6 across vertebrates in stabilizing Dcst1/2. Alternatively, due to Spaca6’s structural similarity to Izumo1 (Nishimura *et al.*, 2016) - a well-known adhesion factor - and the inability of *spaca6* KO sperm to bind to the egg surface in zebrafish, Spaca6 could be involved in sperm-egg adhesion, which may or may not depend on its regulation of Dcst1/2. In this context, studies of its potential interaction partner(s) on the egg membrane in zebrafish and mice might identify new fertilization factors on the egg. Finally, the notion of a membrane complex needed for binding and fusion has been previously proposed (Barbaux *et al.*, 2020; Noda *et al.*, 2020). Spaca6 may contribute to forming and/or stabilizing such a multi-factor complex on the sperm membrane that regulates both binding and fusion. Further investigation of the co-regulation and potential interaction between Spaca6, Dcst1/2, Izumo1 and other known essential fertilization factors may help elucidate the mechanism of gamete fusion on a molecular level.

## Conflict of Interest

The authors declare that the research was conducted in the absence of any commercial or financial relationships that could be construed as a potential conflict of interest.

## Author Contributions

VED and AP conceived the study; MIB, AK and KP designed, performed and analyzed experiments with contributions from VED and AP; AS performed the phylogenetic analysis; VED and AP coordinated and supervised the project; all authors contributed to writing the manuscript.

## Funding

This research was funded by the Research Institute of Molecular Pathology (IMP), Boehringer Ingelheim, the Austrian Academy of Sciences, FFG (Headquarter grant FFG-852936), the FWF START program (Y 1031-B28) to AP, the HFSP Career Development Award (CDA00066/2015) and the HFSP Young Investigator Award to AP, EMBO-YIP funds to AP, and an HFSP post-doctoral fellowship to VED. The funders had no role in study design, data collection and analysis, decision to publish, or preparation of the manuscript. For the purpose of Open Access, the author has applied a CC BY public copyright license to any Author Accepted Manuscript (AAM) version arising from this submission.

## Acknowledgments

We thank the team of the Biooptics facility at the Vienna Biocenter, in particular Pawel Pasierbek and Thomas Lendl, for support with microscopy and image analysis; the animal facility personnel from the IMP for taking excellent care of zebrafish; Anna Bandura for help with genotyping; Juraj Ahel for providing the AlphaFold2 script; Andreas Blaha and Sara Berent for valuable help with the sperm morphology assay; and the entire Pauli lab for fruitful discussions.

## Ethics Statement

All animal experiments were conducted according to Austrian and European guidelines for animal research and approved by the Amt der Wiener Landesregierung, Magistratsabteilung 58 - Wasserrecht (animal protocols GZ: 342445/2016/12 and MA 58-221180-2021-16).

## Data availability statement

The original contributions presented in the study are included in the article/supplementary material; further inquiries can be directed to the corresponding author/s.

## Materials and Methods

### 1.1 Zebrafish husbandry

Zebrafish *(Danio rerio)* were raised according to standard protocols (28°C water temperature, 14/10 hour light/dark cycle). TLAB fish were generated by crossing AB and natural variant TL (Tupfel Longfin) zebrafish and used as wild type for all experiments. The generation of zebrafish *spaca6* KO fish is described below. The *dcst2^−/−^* zebrafish line has been described previously (Noda *et al.*, 2021). All fish experiments were conducted according to Austrian and European guidelines for animal research and approved by the Amt der Wiener Landesregierung, Magistratsabteilung 58 - Wasserrecht.

### 1.2 Identification of the full-length zebrafish Spaca6 sequence

Current gene annotations for zebrafish *spaca6* (NCBI *Danio rerio* Annotation Release 106: XM_021466914.1, 7 exons; and ENSEMBL release 104: BX539313.2-201, ENSDART00000155083.2, 8 exons) were found in the Genome Browser (http://genome.ucsc.edu) using the zebrafish genome release (GRCz11). To obtain the correct, full-length sequence for zebrafish Spaca6, wild-type zebrafish testis cDNA was used for amplifying a region predicted to encompass the full-length protein sequence (primers used for PCR: Spaca6_CDS_F: GCTACTTGTTCTTTTTGCAGGATCCGCCACCATGTTTGTGTTTATTGCAAAAC and Spaca6_CDS_R: ACACTCCTGATCCTCCTGAGAATTCGGCTGGATTAGAAACGTTG). The amplified region was cloned and subsequently sequence-verified. Published RNA sequencing data from adult tissues (Herberg *et al.*, 2018; Noda *et al.*, 2021) was used to analyse *spaca6* gene expression levels in various adult tissues and to examine the coverage tracks across spaca6 in testis and oocytes using the Integrative Genomics Viewer (IGV) (http://software.broadinstitute.org/software/igv/).

### 1.3 Taxonomic tree of Spaca6, Izumo1 and DC-STAMP-like proteins and analysis of Spaca6 protein structure and conservation

Spaca6 orthologs were collected in a series of NCBI blast searches starting with human SPACA6 (sp|W5XKT8|SACA6_HUMAN) and zebrafish Spaca6 (ref|XP_021322589.1|) from the UniProt reference proteomes or NCBI non redundant (nr) protein database applying significant E-value thresholds (1e-05) (Altschul, 1997; Schoch *et al.*, 2020; Bateman *et al.*, 2021). Sequences of the DC-STAMP protein family, including human DCST1, DCST2, DC-STAMP, and OC-STAMP, were identified in a Hidden Markov Model (HMM) search using the PFAM DC-STAMP model against UniProt reference proteomes databases applying significant E-value thresholds (< 0.01) (Eddy, 1998; El-Gebali *et al.*, 2019). Izumo1 protein family members were identified using the PFAM IZUMO HMM search tool, covering the aminoterminal conserved region of human IZUMO1 (21-165, sp|Q8IYV9|IZUM1_HUMAN) in the UniProt reference proteomes databases (E-value < 0.01). In addition, an extended region of IZUMO1 (corresponding to human 1-219) was used and searched for with NCBI blastp in the NCBI nr database (E-value < 0.001). Out of the full set of 453 taxa containing either DC-STAMP, Izumo1 or Spaca6 proteins, 52 representative animal species were selected, and a taxonomic tree was retrieved using the NCBI Taxonomy CommonTree tool (Schoch *et al.*, 2020). The tree visualization was performed in iTOL v6 (Letunic and Bork, 2021).

The protein sequence alignment of vertebrate Spaca6 amino acid sequences was performed using the Muscle alignment tool (http://www.drive5.com/muscle/, version 3.8.31) and visualized with Jalview (Waterhouse *et al.*, 2009).

The protein domain predictions for zebrafish Spaca6 and mouse SPACA6 (Uniprot ID: E9Q8Q8) were obtained from InterPro (Blum *et al.*, 2021). Additionally, the Ig-like domain annotation was derived from previously published data for mouse IZUMO1 and SPACA6 (Nishimura *et al.*, 2016). Secondary and tertiary protein structure predictions were obtained using AlphaFold2 (Jumper *et al.*, 2021). The mouse SPACA6 model (Identifier: AF-E9Q8Q8-F) was already predicted, whilst the zebrafish Spaca6 tertiary structure was modeled using the newly identified Spaca6 amino acid sequence without the signal peptide.

### 1.4 Generation of *spaca6^−/−^* zebrafish

*Spaca6* KO fish were generated by Cas9-mediated mutagenesis. Two guide RNAs (sgRNAs) targeting the 3^rd^ and 4^th^ exons of the full-length *spaca6* gene were generated according to published protocols (Gagnon *et al.*, 2014) by oligo annealing followed by T7 polymerase-driven *in vitro* transcription (gene-specific targeting oligos: spaca6_1_gRNA: TAATACGACTCACTATAGGCGGCCTCAAGCCTGCCCAGGTTTTAGAGCTAGAAATAGCAAG and spaca6_2_gRNA: TAATACGACTCACTATAGGTCTGGATGTTTGCCCCCATGGTTTTAGAGCTAGAAATAGCAAG; common tracer oligo AAAAGCACCGACTCGGTGCCACTTTTTCAAGTTGATAACGGACTAGCCTTATTTTAACTTGCTATTTCT AGCTCTAAAAC). Cas9 protein and *spaca6* sgRNAs were co-injected into the cell of one-cell stage TLAB embryos. Putative founder fish were outcrossed to TLAB wild-type fish. A founder fish carrying a germline mutation in the *spaca6* gene was identified by a size difference in the *spaca6* PCR amplicon in a pool of embryo progeny (primers: spaca6_gt_F: GCAGAGAAATCTTGATTGGAGG and spaca6_gt_R: AAGCAGACCAGTATACAATTTTTGC). Embryos from this founder fish were raised to adulthood. Amplicon sequencing of adult fin-clips identified the 86-nt deletion, which results in a frameshift mutation and a premature stop codon in intron 2 (GRCz11: Chr16:24,907,548). Genotyping of *spaca6* mutant fish was performed by PCR using primers: spaca6_gt_F and spaca6_gt_R. Detection of the deletion was performed by standard gel electrophoresis using a 4% agarose gel. Homozygous *spaca6^−/−^* fish were generated by outcrossing *spaca6^+/−^* fish to wild-type fish and then incrossing *spaca6^+/−^* fish from the next generation.

### 1.5 Generation of zebrafish expressing transgenic Spaca6

The full-length *spaca6* coding sequence, including the *spaca6* signal peptide, the extracellular region and transmembrane and intracellular domains, was amplified by PCR from cDNA derived from adult zebrafish testis (Spaca6_CDS_F and Spaca6_CDS_R) and subcloned by Gibson cloning (Gibson *et al.*, 2009) into a vector for Tol2-mediated transgenesis along with a *SG-linker-T2A-sfGFP* sequence inserted in frame immediately before the stop-codon of the *spaca6* sequence (resulting vector: pMTB Tol2 - *actb2-*promoter - *spaca6* - *SG-linker-T2A-sfGFP* SV40UTR). Zebrafish lines expressing transgenic Spaca6 were generated by injecting the *spaca6* expression construct with *Tol2* mRNA into *spaca6^+/−^* zebrafish embryos (15 ng/μl of the plasmid in RNase-free water, 35 ng/μl *Tol2* mRNA, 0.083% phenol red solution [Sigma-Aldrich]), following standard procedures. Injected embryos with high expression of sfGFP at one day post fertilization were raised to adulthood. Putative founder fish were crossed to *spaca6^−/−^* or *spaca6^+/−^* fish and the progeny was screened for fluorescence, raised to adulthood and genotyped using primers spaca6_gt_F and spaca6_gt_R to identify adult fish lacking endogenous *spaca6. spaca6^−/−^* male fish expressing transgenic *spaca6 (spaca6^−/−^; tg(spaca6))* were crossed to wild-type females in order to quantify fertilization rates and assess functionality of the construct.

### 1.6 Quantification of *in vivo* fertilization rates in zebrafish

The evening prior to mating, male and female fish were separated in breeding cages (one male and one female per cage). The next morning, male and female fish were allowed to mate. Eggs were collected and kept at 28°C in E3 medium (5 mM NaCl, 0.17 mM KCl, 0.33 mM CaCl_2_, 0.33 mM MgSO_4_, 0.00001% Methylene blue). The rate of fertilization was assessed approximately 3 hours post laying. By this time, fertilized embryos have developed to ~1000-cell stage embryos, while unfertilized eggs resemble one-cell stage embryos. Direct comparisons were made between siblings of different genotypes (wild type; *spaca6^−/−^; spaca6^+/−^: spaca6^−/−^ tg(actb2::spaca6-t2a-GFP)).*

### 1.7 Assessment of the sperm morphology

To collect wild-type or mutant sperm, male zebrafish were anesthetized using 0.1% tricaine. Sperm was collected with a glass capillary from the urogenital opening and immediately fixed with 3.7% formaldehyde at 4°C for 20 minutes. Sperm were spun onto an adhesive slide using a CytoSpin 4 (Thermo Fisher Scientific) at 1,000 rpm for 5 minutes followed by permeabilization with ice-cold methanol for 5 minutes and a wash with 0.1% Tween in 1x PBS (PBST). After mounting using VECTASHIELD Antifade with DAPI (Vector Laboratories), sperm were imaged with an Axio Imager.Z2 microscope (Zeiss) with an oil immersion 63x/1.4 Plan-Apochromat DIC objective.

### 1.8 Assessment of sperm motility

Sperm were isolated from 1-2 wild-type and mutant male fish and kept in 100 μl Hank’s saline (0.137 M NaCl, 5.4 mM KCl, 0.25 mM Na_2_HPO_4_, 1.3 mM CaCl_2_, 1 mM MgSO_4_, and 4.2 mM NaHCO_3_) containing 0.5 μM MitoTracker Deep Red FM (Molecular Probes) for >10 minutes on ice. Sperm (approximately 5000 sperm/μl) were activated using E3 medium in a 1:4 dilution and placed onto a 10 μm thick chamber slide (Leja counting chamber, SC 10-01-04-B). Sperm motility was imaged 30 seconds after activation using an Axio Imager.Z2 microscope (Zeiss) and a 10x/0.3 plan-neofluar objective using darkfield. Sperm tracks were analyzed using Fiji with the “manual tracking” plugin (Cordelieres, 2005). Sperm that were present in the movie for more than 30 timeframes were tracked for as many frames as possible. Coordinates of the sperm cells were used to calculate average sperm speed and displacement. Sperm displacement was calculated by measuring the distance between the first and the last coordinates (normalized by 100 timeframes).

### 1.9 Imaging of zebrafish sperm approach

Sperm were isolated from 2-4 wild-type and mutant male fish and kept in 150 μl Hank’s saline containing 0.5 μM MitoTracker Deep Red FM (Molecular Probes) for >10 minutes on ice. Unactivated, mature eggs were then isolated from a wild-type female. To prevent activation, eggs were kept in sorting medium (Leibovitz’s medium, 0.5% BSA, pH 9.0) at room temperature. Eggs were kept in place using a petri dish with cone-shaped agarose molds (1.5% agarose in sorting medium) filled with sorting medium. Imaging was performed with a LSM800 Examiner Z1 upright system (Zeiss) with a 20x/1.0 plan-apochromat water dipping objective. Before sperm addition, sorting media was removed and 1 ml of E3 medium carefully added close to the egg. 3 μl of stained spermatozoa (approximately 150,000 – 300,000 sperm) was added as close to the egg as possible during image acquisition. The resulting time-lapse movies were analyzed using Fiji. Timestamps were calculated beginning with the addition of sperm to the eggs.

### 1.10 Imaging and analysis of zebrafish sperm-egg binding

Sperm were isolated from 2-4 wild-type and mutant male fish and kept in 200 μl Hank’s saline containing 0.5 μM MitoTracker Deep Red FM (Molecular Probes) on ice. Unactivated, mature eggs were squeezed from a wild-type female fish and activated by addition of E3 medium. After 10 minutes, 1-2 eggs were manually dechorionated using forceps and 1 egg was transferred to a cone-shaped 2% agarose-coated imaging dish with E3 medium. After focusing on the egg plasma membrane, the objective was briefly lifted to add 2-5 μl of stained sperm (approximately 200,000 - 250,000 sperm). Imaging was performed with a LSM800 Examiner Z1 upright system (Zeiss) using a 20x/0.3 Achroplan water dipping objective. Images were acquired until sperm were no longer motile (5 minutes). To analyze sperm-egg binding, stably bound sperm were counted. Sperm were counted as bound when they remained in the same position for at least 2 minutes following a 90-second period in which they activated and approached the egg. Data was plotted as the number of sperm bound per 100 μm of egg membrane for one minute.

### 1.11 Sperm immunocytochemistry

Sperm from zebrafish males was collected in 3.7% formaldehyde diluted in Hank’s saline solution and stored on ice for 20 minutes to 1 hour. Sperm was pelleted by centrifugation at 850 rpm for 3 minutes, and the fixative was replaced with Hank’s saline. Sperm was spun onto an adhesive slide with a CytoSpin 4 (Thermo Fisher Scientific) at 800 rpm for 3 minutes. Slides were briefly washed once in 1x PBS, and the sperm was permeabilized in 0.25% Tween in 1x PBS for 30 minutes before blocking with 10% normal goat serum (Invitrogen) and 40 μg/ml BSA in PBST for at least 1 hour. Slides were then incubated with mouse anti-zebrafish-Dcst2 antibody in blocking buffer (1:650; (Noda *et al.*, 2021)) overnight at 4°C in a humidified chamber. After several washes with PBST, slides were incubated with goat anti-mouse IgG Alexa Fluor 488 secondary antibody (1:380, Thermo Fisher Scientific) for 1 hour, washed several times with PBST and finally once with 1x PBS. After mounting using VECTASHIELD Antifade with DAPI (Vector Laboratories), sperm was imaged with an Axio Imager.Z2 microscope (Zeiss) using an oil immersion 100x/1.4 plan-apochromat objective. Widefield sperm images were processed for each genotype using Fiji by adjusting image brightness and contrast without clipping of intensity values.

### 1.12 Western blotting of sperm samples

For western blot analysis, sperm from 3-6 males was sedimented at 3000 rpm for 3.5 minutes. The supernatant was replaced with RIPA buffer (50 mM Tris-HCl [pH 7.5], 150 mM NaCl, 1 mM MgCl_2_, 1% NP-40, 0.5% sodium deoxycholate, 1x complete protease inhibitor [Roche]) including 1% SDS. After preparation of all samples, 1 U/μl benzonase (Merck) was added, and samples were incubated for 30 minutes at room temperature. Samples were then mixed with 4x Laemmli buffer containing β-mercaptoethanol and boiled at 95°C for 5 minutes. After SDS-PAGE, samples were wet-transferred onto a nitrocellulose membrane. Total protein was visualized by Ponceau staining before blocking with 5% milk powder in 0.1% Tween in 1x TBS (TBST). Membranes were incubated in primary mouse anti-zebrafish-Dcst2 antibody (1:500; (Noda *et al.*, 2021)) overnight at 4°C, then washed with TBST before HRP-conjugated secondary antibody (1:10.000 [115-036-062, Dianova]) incubation for 1 hour. Membranes were washed several times in TBST before HRP activity was visualized using Clarity Western ECL Substrate (BioRad) on a ChemiDoc (BioRad). For visualizing Tubulin levels, membranes were stripped using Restore Western Blot Stripping Buffer (Thermo Fisher Scientific) before washing, blocking and incubation with mouse anti-alpha-Tubulin antibody (1:20.000 [T6074, Merck]) and proceeding with secondary antibody staining and detection as described above. To assess relative Dcst2 protein amounts, average intensities of Dcst2-specific bands were quantified in Fiji on 3 independent immunoblots for each genotype relative to tubulin levels, which was used as loading control. Values were then normalized to the levels of wild-type sperm.

### 1.13 Data visualization and statistical analyses

Data plotting and statistical analyses were performed in GraphPad Prism. Wherever possible, data was plotted as individual data points and indicating the standard deviation (SD).

## Supplementary Information

### Supplementary Figures

**Supplementary Figure S1.**
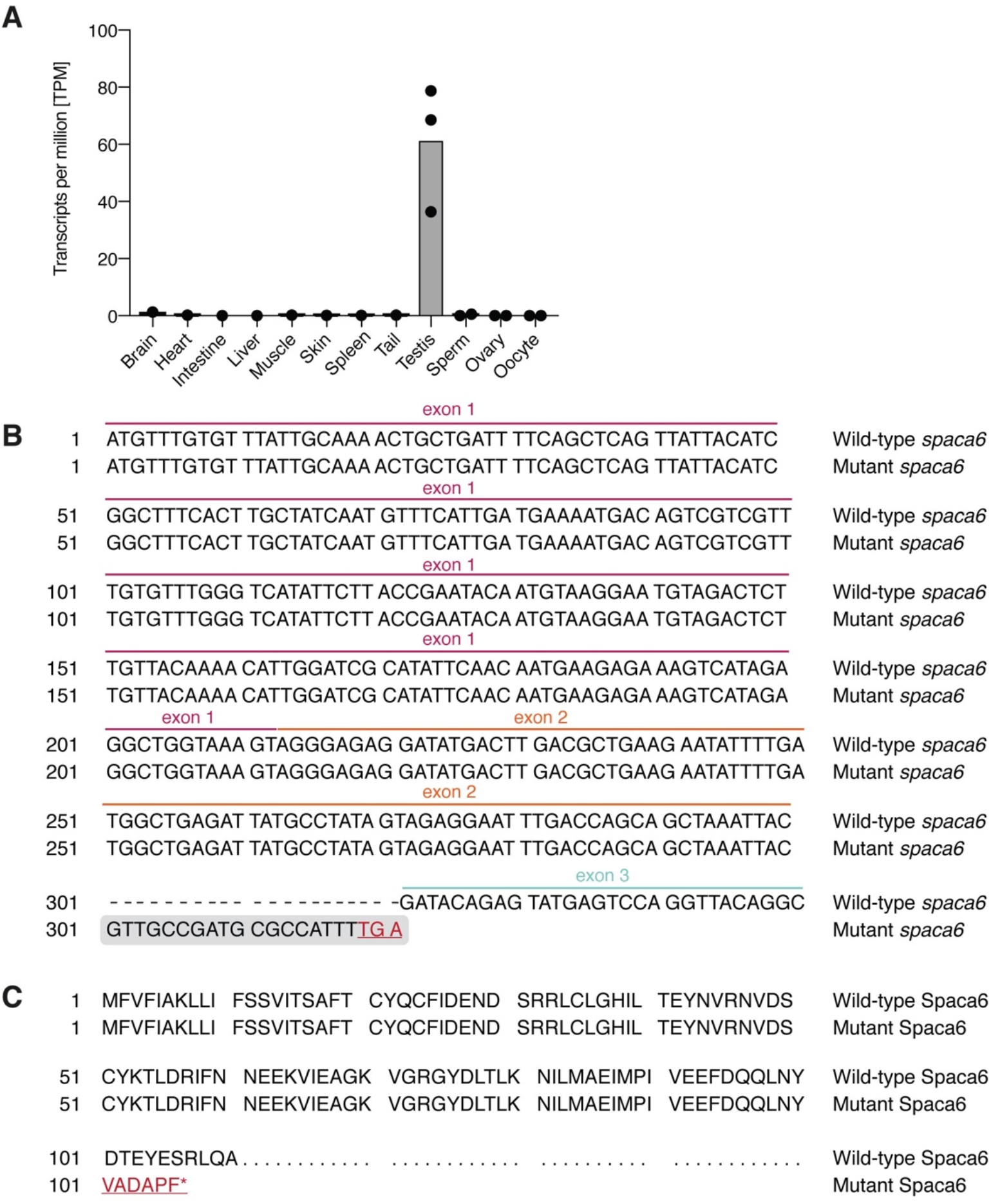
Characterization of *spaca6* in zebrafish. **A.** RNA-Seq analysis of *spaca6* gene expression levels in various adult tissues. Gray bar, mean. TPM, transcripts per million. **B.** Partial cDNA sequence alignment of wild-type and mutant *spaca6.* The first 329 base pairs of the wild-type cDNA sequence were aligned to the mutant cDNA sequence, which retains part of intron 2 (shaded in gray) and contains a stop codon (red, underlined). **C.** Partial amino acid sequence alignment for wild-type (amino acids 1-110) and mutant (full-length) Spaca6. Retained intronic sequence in mutant *spaca6* leads to the translation of 6 additional amino acids before a premature stop codon (red, underlined).

**Supplementary Figure S2.**
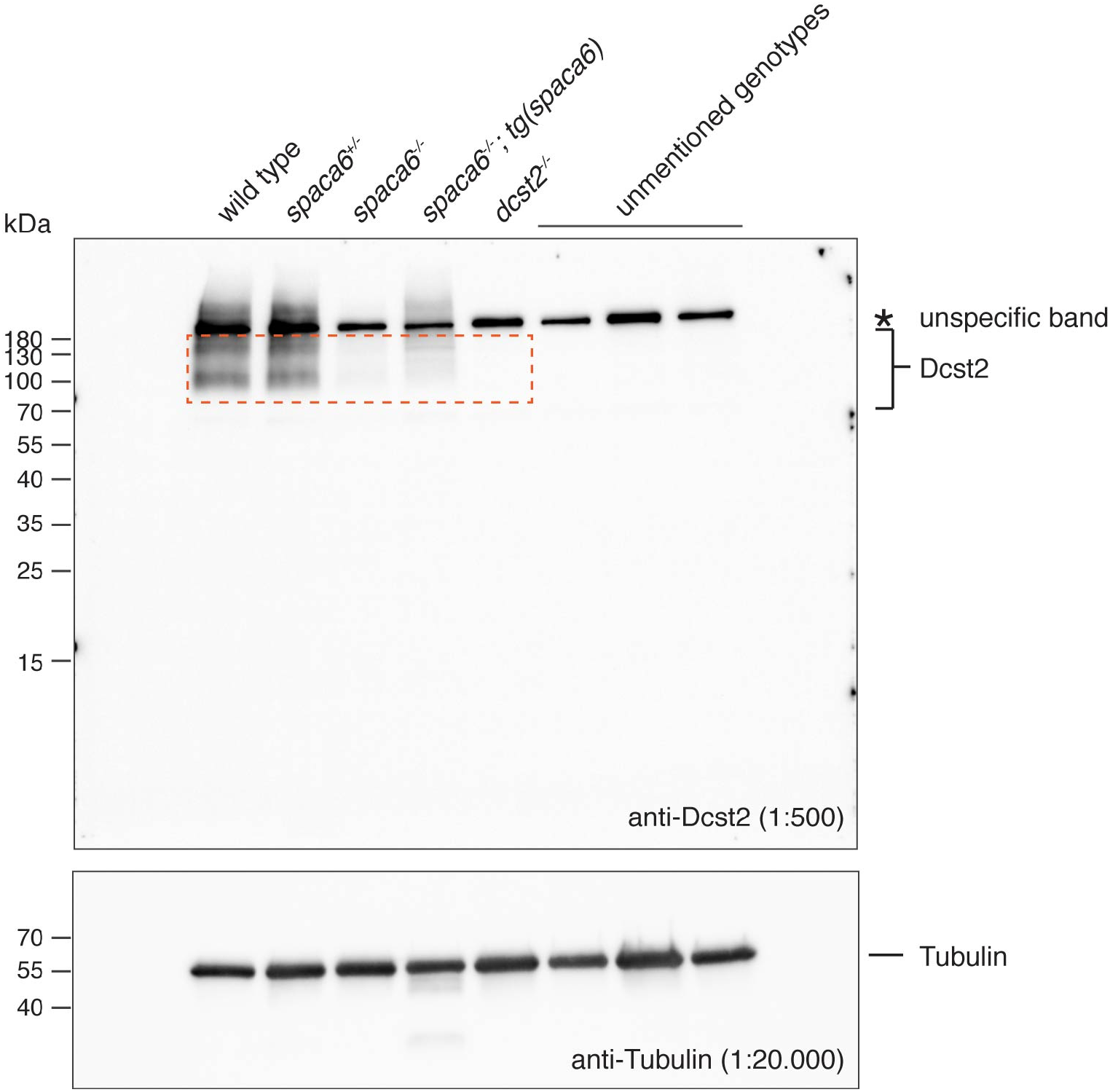
Dcst2 levels are reduced in zebrafish *spaca6* KO sperm. Uncropped immunoblot of sperm samples probed with antibodies against zebrafish Dcst2. An unspecific band (asterisk) is detected in all genotypes above 180 kDa. The cropped region is highlighted by a red-dashed rectangle.

### Supplementary Movies

**Movie 1: Wild-type and *spaca6^−/−^* sperm can approach the micropyle.** Wild-type or *spaca6^−/−^* sperm stained with MitoTracker Deep Red (red) were added to wild-type eggs and imaged following sperm addition. Scale bar = 50 μm.

**Movie 2: *Spaca6^−/−^* sperm are unable to stably bind to wild-type eggs**. Wild-type or *spaca6^−/−^* sperm stained with MitoTracker Deep Red (red) were added to dechorionated wild-type eggs and imaged following sperm addition. Two minutes after sperm addition, wild-type sperm are stably bound to the egg membrane while *spaca6^−/−^* sperm are unable to bind. Scale bar = 50 μm.

## Notes

### Competing Interest Statement

The authors have declared no competing interest.

